# Leveraging Deep Learning and MD Simulations to Decipher the Molecular Basis of Attenuated Activity in Glycocin F

**DOI:** 10.1101/2025.10.01.679699

**Authors:** Dheeraj Kumar Sarkar, Subinoy Adhikari, Avadhesha Surolia

## Abstract

The escalating crisis of multi-drug resistant bacteria demand a new generation of antibiotics. Glycocin F (GccF), a potent bacteriocin, is a promising candidate, but its function hinges on unique post-translational glycosylation. Intriguingly, a seemingly minor chemical tweak of α-methylation at Ser18 of GccF destroys its activity, reducing potency by 1000-fold. To quantify how this subtle chemical change leads to profound functional compromise, we used an advanced molecular dynamics framework guided by Variational Autoencoder to unravel GccF’s complex dynamics. Our findings reveal that native glycosylation preserves conformational plasticity to maintain functionally relevant conformations. In stark contrast, α-methylation introduces local rigidity, locking the peptide into fewer metastable basins with significantly slower transition rates. This leads to the disruptions of α-helix structure which perturbs the loop-helix coupling and trapping the peptide into non-functional conformations. Together, these findings provide a critical blueprint for the rational design of next generation antibiotics, demonstrating how precise chemical modifications can dictate a peptide’s function by profoundly altering its structural dynamics.

## 1 Introduction

The emergence of multi-drug resistance bacteria constitute a major challenge to global health. One such example is vancomycin-resistant *Enterococcus* species (VRE), which have rendered many traditional antibiotics ineffective [1-2]. Thereby motivating exploration of new antimicrobials agents with novel mechanisms of action, among those bacteriocins, ribosomally synthesized and post-translationally modified peptides (RiPPs) hold a considerable promise [3-4].

Secreted by *Lactobacillus plantarum* KW30, GccF stands out as a particularly compelling candidate within this class. GccF is a 43-residue bacteriocin that demonstrates potent and reversible bacteriostatic activity against a range of Gram-positive bacteria at remarkably low picomolar concentrations [5-6]. Beyond its disulfide connectivity (Cys-5 to Cys-28 and Cys-12 to Cys-21), GccF is distinguished by its post-translational modifications (PTMs), specifically its diglycosylated state which is obligatoty for the function of the protein. The first is a β-O-linked GlcNAc at Ser18, which is situated on the interhelical loop of the peptide [5-7]. The second, and more unusual, modification is a β-S-linked GlcNAc at the C-terminal Cys43 [5-6]. The removal of this single sugar moiety completely abolishes the peptide’s function, suggesting a primary and indispensable role [6,8].

As a general post-translational modification, glycosylation is well-known to confer enhanced biophysical stability to peptides and proteins. This includes a significant increase in both thermal stability and resistance to proteolytic degradation, which are critical properties for the viability of any therapeutic peptide [9-13]. A recent study revealed an instructive counterexample to its general utility by application of α-methylation to GccF at Ser18. While the modification is typically used to rigidify peptides and increase their stability, the study revealed substitution of the Cα proton with a methyl group at Ser18 resulted in a catastrophic loss of antimicrobial activity, reducing its potency by an astonishing 1000-fold [6] and hence showing a marked difference in the orientation and mobility of the critical interhelical loop.

Since, the study by Harjes *et al*., 2023 [6], argued the structural basis of this functional loss by GccF by a subtle chemical change, we here used a framework to investigate beyond the static structural comparison in order to quantify how this modifications leads to a profound functional loss. We analyzed three molecular systems: control, glycosylated and methylated to describe a non-glycosylated control, the native glycosylated peptide (Ser18-GlcNAc, Cys43-GlcNAc), and an α-methyl-Ser18 glycosylated construct respectively. All-atom molecular dynamics (MD) simulations are the ideal starting point, as they provide atomic-level trajectories of biomolecular systems over time [14]. In order to interpret the high-dimensional trajectory data from unbiased MD of aggregated 3 µs data for each molecular systems, we employed a powerful integrative pipeline that leverages machine learning with Variational Autoencoder (VAE) for dimensionality reduction [15-17]. A key innovation in this framework is the use of a tailored β-annealing schedule for VAE training as standard VAE often struggles with “posterior collapse,” where the model fails to learn a meaningful latent representation [18]. By dynamically adjusting the β factor, we reconstructed a more robust latent representation of our projected trajectory data [16]. The refined latent space from the VAE is then used as the basis for building Markov State Models (MSMs). MSMs are a statistical mechanics tool for extracting kinetic information from MD trajectories [19-20]. The combination of all-atom MD combined with β-annealing VAEs guided by MSMs provided an ensemble-level view of GccF’s conformational dynamics, allowing for a precise quantification of how glycosylation and α-methylation modulating the dynamics and stability of the critical interhelical loop in GccF. We observed α-methylation may have locally rigidified the Ser18 residue, this local constraint appears to have propagated a global conformational effects. This in turn, compromised the peptide’s ability to adopt the native, functional ensemble required for its activity, suggesting that the precise positioning and presentation of the glycan moiety is critical for its function. The results will not only elucidate the precise mechanism of action for this unique antimicrobial peptide but will also provide broader insights into the sophisticated role of PTMs in dictating peptide function and stability, the knowledge of which will pave the rational design of future generation of antibiotics.

## 2 Methods

### 2.1 Molecular Dynamics Simulations

Molecular dynamics (MD) simulations were conducted using GROMACS version 2021.6 [21] with the CHARMM36 force field [22-23]. Three variants of GccF were modeled: (i) a *control* system without glycosylation, where both GlcNAc moieties were removed; (ii) a *glycosylated* system based on the native NMR structure (PDB ID: 2KUY) [24]; and (iii) a *methylated* system prepared using the NMR structure (PDB ID: 8DFZ) [6]. Bonded and non-bonded parameters for the α-methylated Ser18 linked GlcNAc residue (SI Figure S1) were calculated using the DFT Seminario method [25] within the Gaussian 16 software package [26] to derive force constants, which were used for all-atom MD simulations. Disulfide bonds between Cys4–Cys39 and Cys9–Cys43 were enforced in all systems.

Each system was solvated in a cubic box of TIP3P water [27-28], extending at least 1.0 nm beyond the peptide surface, and neutralized with Na^+^/Cl^−^ ions to achieve a physiological ionic strength of 150 mM. Systems were energy-minimized using the steepest-descent algorithm until the maximum force was below 100 kJ mol^−1^ nm^−1^. Equilibration was performed in two stages: (i) 2 ns of NVT equilibration using the V-rescale thermostat [29] with position restraints on heavy atoms, and (ii) 2 ns of NPT equilibration using the Parrinello–Rahman barostat [30]. Production MD simulations were conducted in the NPT ensemble with coordinates saved every 100 ps. All bonds were constrained using the LINCS algorithm [31], and long-range electrostatics were treated with the Particle Mesh Ewald (PME) method [32] using a 1.2 nm cutoff for both Coulomb and van der Waals interactions. Each system was simulated in triplicate to ensure reproducibility.

### 2.2 Variational Autoencoder and Free-energy Surface Construction

Molecular dynamics trajectories for the *control, glycosylated*, and *methylated* systems were processed to remove solvent molecules and ions, and periodic boundary conditions were corrected, to extract features for dimensionality reduction. For each trajectory, the protein backbone dihedral angles, specifically the *Φ* and *Ψ* torsion angles, were computed. To properly represent the circular nature of this angular data for the neural network, the cosine and sine of each angle were calculated, resulting in four feature components for each residue. These components were combined into a single feature vector for each frame, and the complete dataset for each system was subsequently partitioned into training and test sets using a 90:10 split. A β-Variational Autoencoder (VAE) [16] was employed to construct a non-linear, low-dimensional representation of the dihedral angle space. The encoder network consisted of three fully connected layers with 168, 64, and 16 neurons, mapping into a two-dimensional latent space.

The decoder network was symmetrically structured, with layers of 16, 64, and 168 neurons. Training was performed for 1000 epochs using Stochastic Gradient Descent with a learning rate of 0.001, with batch sizes of 2048 for training and 2000 for validation. The total loss function was defined as

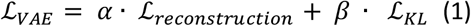

where the reconstruction loss was given by the mean squared error (MSE), ℒ_*KL*_ denotes the Kullback–Leibler divergence, α was fixed at 10, and β was annealed during training.

The β parameter was initially annealed from 0 to 1 during training using linear, cosine, and logistic schedules. For each system, the optimal β value was determined as the one that maximized the fraction of variance explained (FVE), following the procedure established in prior work. The VAEs were subsequently retrained by annealing β to these system-specific optimal values. Model selection among the annealing schemes was carried out using 5-fold cross-validated VAMP-2 scores (SI Figure S6). Among the tested approaches, cosine annealing VAE consistently yielded the highest scores, surpassing both the alternative annealing schedules and the leading two dimensions obtained via Principal Component Analysis (PCA). We investigated the free energy surface (FES) profile by plotting the first two principal components from PCA and projecting the trajectory data using the dihedral matrix into a lower dimensional space using our VAE approach. This indicated the efficiency of VAE over traditional PCA in capturing non-linear, collective variables that are crucial for describing complex protein dynamics (SI Figure S5 & S12).

### 2.3 Markov State Model (MSM) Analysis and Validation

We tested the convergence of all simulated MD trajectories of 3 µs each for *control, glycosylated* and *methylated* systems using time-dependent behavior and auto-correlation functions analysis [33]. We analyzed the change over time and the autocorrelation of two key metrics: the *Cα end-to-end distance* (the distance between the Cα atoms of the first and last residues) and the *Cα RMSD* (the root-mean-square deviation of each simulation frame from the initial frame). A convergence within 0.5 µs can be observed for all the three replicates (SI Figure S2) generated for the three molecular systems. The choice of the cosine-annealed VAE is supported by model selection: across annealing schemes and PCA baselines. Consequently, the cosine-annealed VAE, with optimal β values of 0.085 *(control)*, 0.135 *(glycosylated)*, and 0.024 *(methylated)* were chosen to generate the two-dimensional latent spaces used for the construction of the MSMs and cosine annealing yielded the highest VAMP-2 score (SI Figure S5-6). All MSMs were generated using PyEMMA package [34]. The slow processes evolved from the MD trajectories were analyzed using implied timescale (ITS) plot (SI Figure S8) and the number of macrostates were chosen based on spectral analysis (SI Figure S9).

#### 2.3.1 Analysis and Validation

The constructed MSMs were validated using Chapman-Kolmogorov [35] test (SI Figure 9-11) against estimated models until lag time of 100ps. All trajectories were analyzed using MDAnalysis [36] and custom Python scripts. Backbone root-mean-square deviation (RMSD) and per-residue root-mean-square fluctuation (RMSF) were calculated relative to the initial conformations. Secondary structure content was determined using the DSSP algorithm [36]. The S^2^ order parameters were computed using the cpptraj [38] module (version 6.18.1) from the AMBER software suite [39], and the backbone NH bond vectors were used to determine the S^2^ values after removing the global rotational and translation motion using GROMACS v2021.6 [21].

## 3 Results

Using our MD-VAE-MSM framework, we studied the role of glycosylation in wild-type GccF *(glycosylated)*, modified Ser18 GccF *(methylated)* along with *control* dataset with no glycosylation. We simulated each molecular systems (Figure 1A) for an aggregated ∼3 µs consisting three replicate runs to study the distinct impacts of glycosylation and α-methylation on GccF’s conformational landscape and functional relationship. β-annealed VAE (Figure 1B) was able to explore a wider conformational minima for the three molecular systems (Figure S5 & 12).

**Figure 1.**
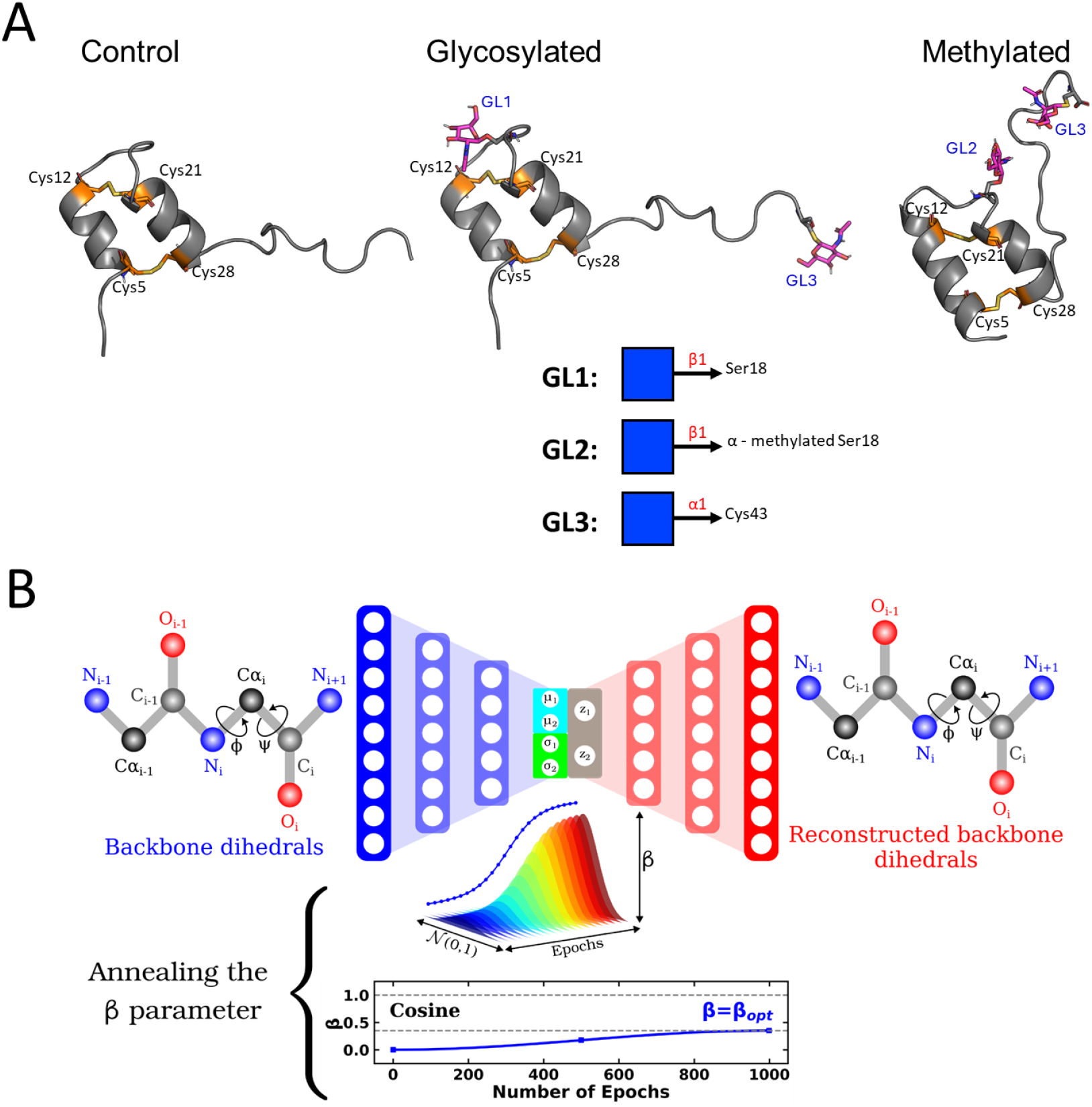
A. Structural representation of Glycocin F (GccF) showing the two glycosylation sites at Ser18 (β-O-GlcNAc) and Cys43 (β-S-GlcNAc), stabilized by nested disulfide bonds (Cys4–Cys39 and Cys9–Cys43). B. Schematic of β-annealed Variational AutoEncoder. A schematic of the reconstructed dihedrals are shown along with the β optimal (β_opt_) annealed using cosine method.

### 3.1 Stability and flexibility profiles: glycosylation and α-methylation stabilizes key loop dynamics

Backbone RMSD profiles revealed that α-methylation promoted overall conformational stability to GccF structure followed by glycosylation with >2.0 Å on average compared to ∼2.5 Å in the control (Figure 1A). RMSF analysis pinpointed reduced fluctuations and prominent rigidity in the interhelical loop (residues 10–19) upon O-glycosylation in wild-type GccF and modified GccF, whereas α-methylation at Ser18 introduced enhanced rigidity in this region (Figure 2B & 2D). These findings align with previous reports linking O-GlcNAc modification to enhanced peptide rigidity and resistance to unfolding [6]. MSM results revealed the transition time from “closed” to “open” state is much higher for the *glycosylated* and *methylated* models (Figure 2C) compared to *control* dynamics depicting much improved and effective role of both O-/S-linked GlcNAc and α-methylation for the stabilization of the scaffold, with the largest effects concentrated in the interhelical loop residues 10-19, that are poised to modulate glycan presentation. S^2^ order analysis (Figure 2B & 2D) revealed Ser18 is the most responsive region which is in good agreement with the root mean square fluctuation (RMSF) analysis. Overall, the α-methylated variant further rigidifies the Ser18-proximal interhelical loop.

**Figure 2.**
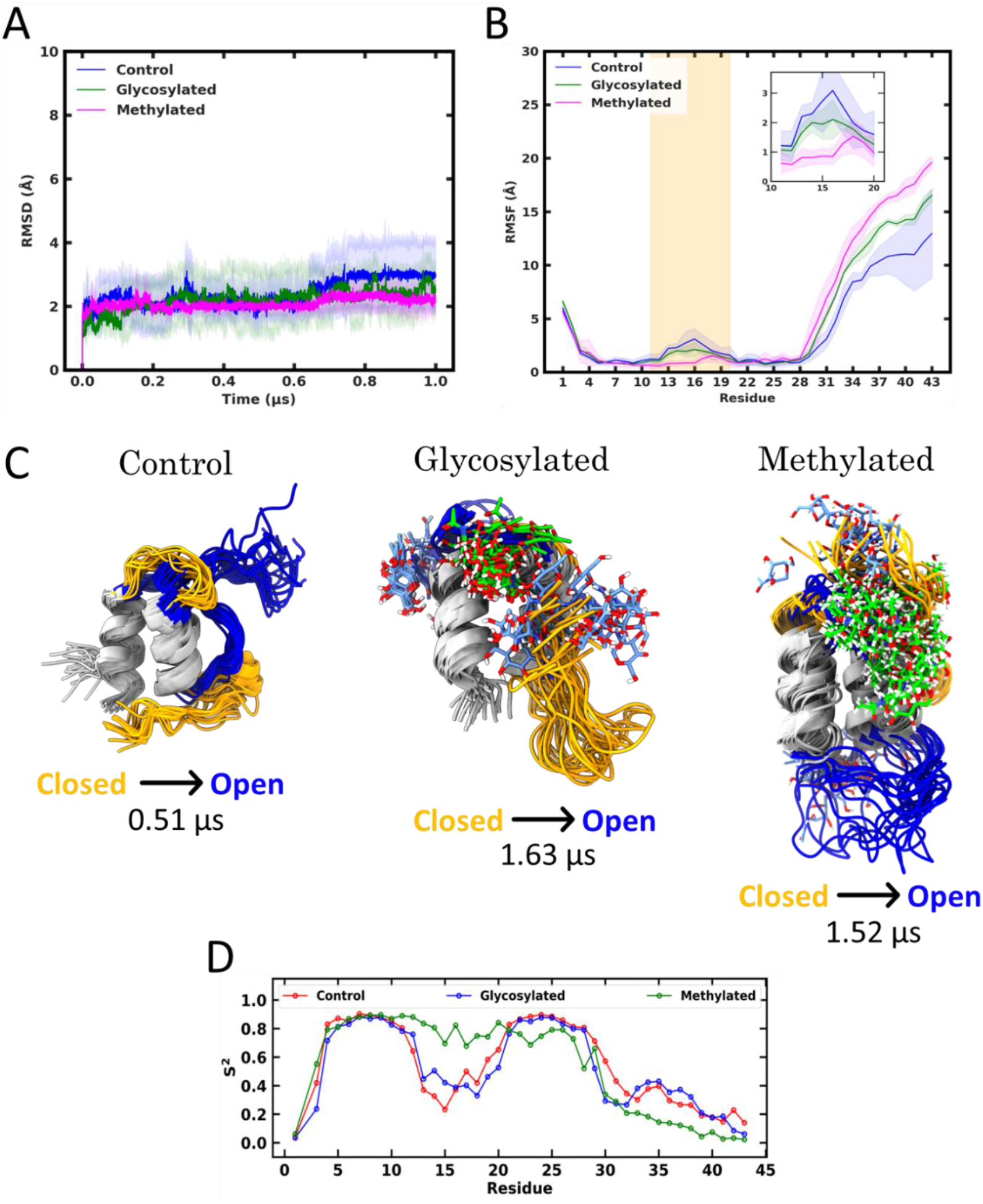
A. Backbone RMSD and B. per-residue RMSF comparisons among *control, glycosylated*, and *methylated* systems. C. Transition time of closed to opened metastable states obtained from MSM analysis for the three molecular systems. D. S^2^ order analysis for *control, glycosylated*, and *methylated* systems.

Secondary structure analyses (SI Figure S3) confirmed that glycosylation preserved helical segments, in contrast to the *methylated* system which exhibited partial loss of α-helical content. Hydrogen-bond analyses (SI Figure S4) revealed prominent stabilization effect by glycan-peptide contacts in the *glycosylated* system, reinforcing glycan modulated loop dynamics and stabilizing the disulfide-anchored scaffold. In contrast, Ser18 α-methylation in *methylated* model showed less prominent H-bonds compared to *glycosylated* model that leads to disrupted H-bonding patterns in helical and interhelical loop regions due to steric constraints facilitated by the O-/S-linked GlcNAc moiety. This altered backbone dihedral preference could result in reduced stabilizing effect of glycosylation for the *methylated* model.

### 3.2 Kinetic decomposition and metastable states in the glycosylated and α-methylated GccF compared to control

Mapping backbone dihedrals into φ–ψ space of Ramachandran plot reveals a more clear, modification-dependent topology (Figure 3). The deglycosylated *control* samples most residues in favored α/β regions and generated a diffuse Ramachandran landscape characteristic of a flexible and dynamically unrestrained peptide backbone (Figure 3A). For the *glycosylated* model, the φ–ψ distributions collapse into discrete states located within canonical α-helical (φ ≈ –65°, ψ ≈ –35°) and β-turn regions (φ ≈ –75°, ψ ≈ 140°), indicating more dynamic conformational ensemble, as glycan-protein contacts could stabilize turns-like conformations (Figure 3B). Moreover, the reduced dispersion of dihedral angles demonstrates that glycan-peptide h-bonding (Figure 3B & SI Figure S4) could stabilize specific backbone orientations, particularly around Ser18 and the inter-helical loop. In contrast, the α-methyl-Ser18 variant shows a broader yet stabilized landscape, with density centered on loop-rigidified conformers. α-methylation at Ser18 could add local steric bulk and subtly shift φ–ψ preferences, with a mild redistribution toward the β/PPII sector (ψ ≈ 120–150°) in exposed segment (Figure 3C).

**Figure 3.**
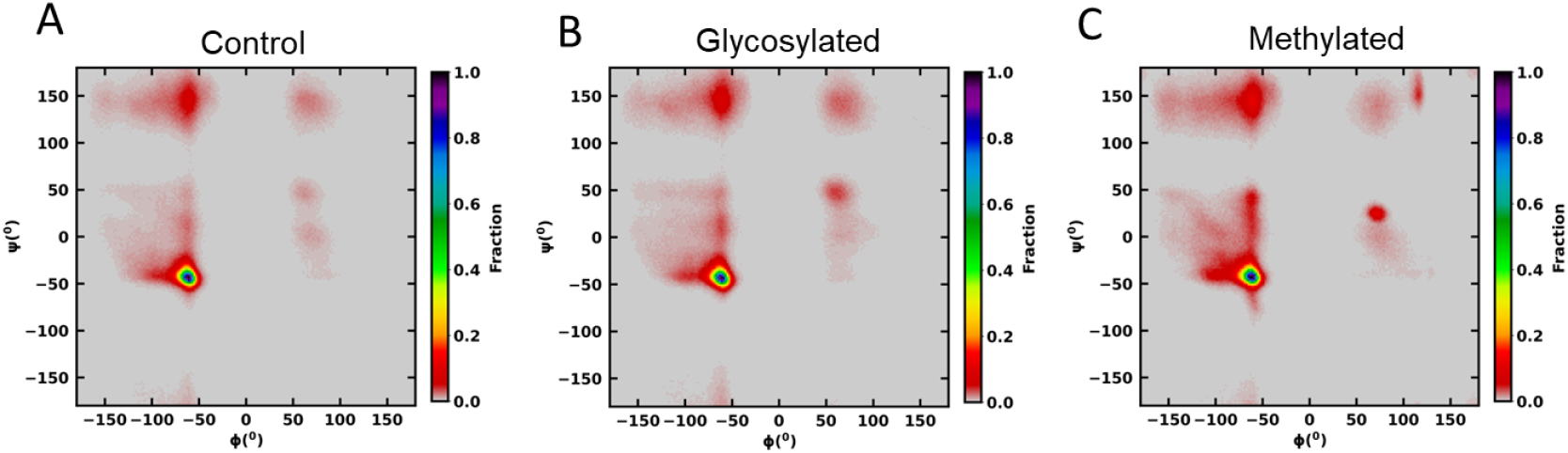
Ramachandran plot comparing backbone conformations of GccF models for A. *control*, B. *glycosylated* and C. *methylated* molecular systems.

Free energy surface (FES) derived from β-annealed VAE latent coordinates revealed that *glycosylated* model resolved into multiple rapidly interconverting states (Figure 4A, SI Tables S1 & SI Figure 12B), while methylation compressed the conformational ensemble into fewer metastable states with much slower transition rated from “closed” to “open” metastates (Figure 5A, SI Table S1 & SI Figure 12C). Consistent with these site-localized dynamics, state-resolved intra-residue contact maps from MSM analyses reveal tighter packing in the stabilized variants compared with the *control*. In the *glycosylated* ensemble, contacts around the Ser18 loop are enriched relative to the *control*, in line with the reduced RMSF observed in Figure 2B,4B & SI Figure S14.

**Figure 4.**
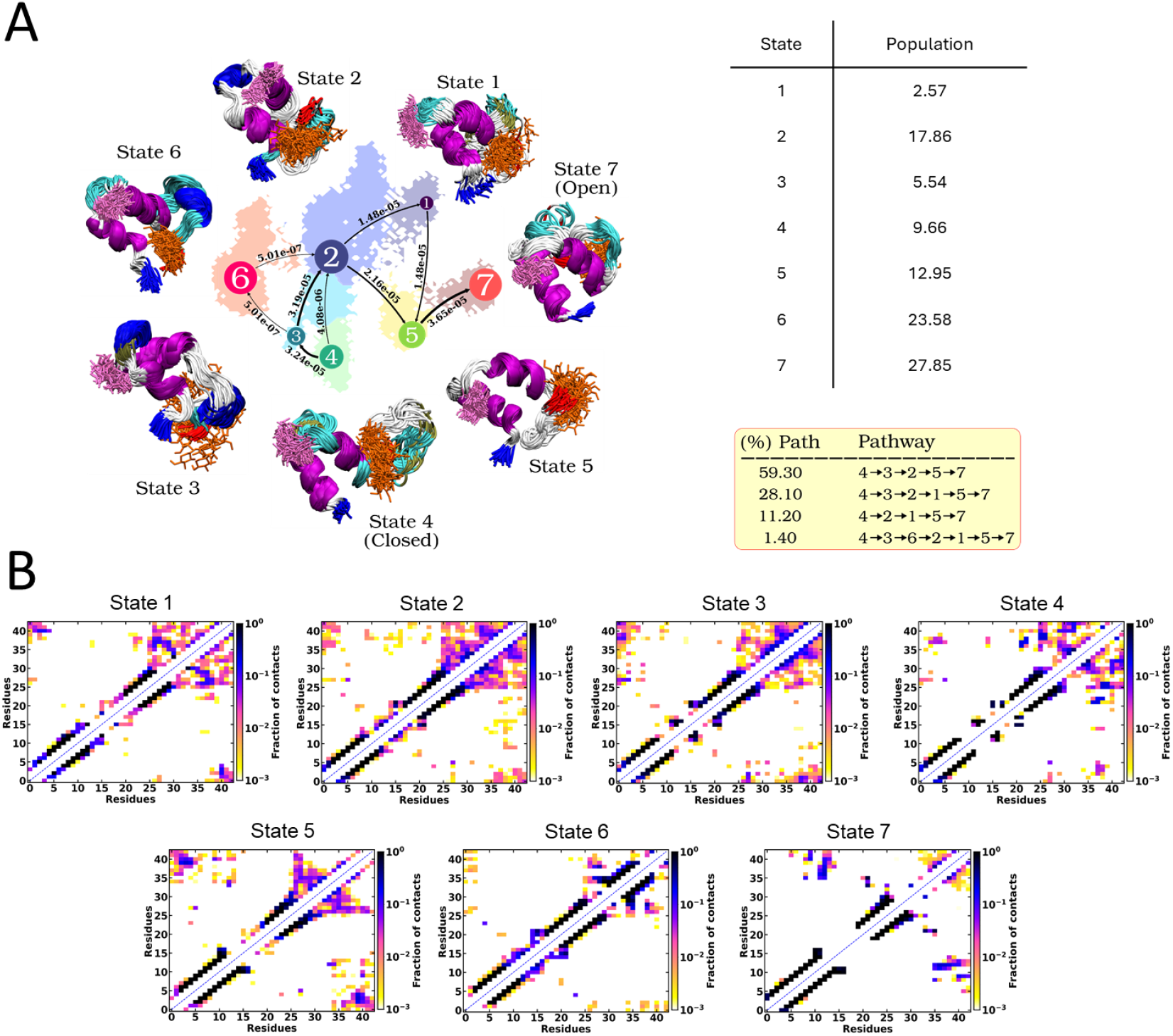
Markov state models and the proportion of each state with unidirectional transition probability and pathways between states derived using latent space of Cosine Annealed VAE from glycosylated dataset. The N-terminal and C-terminal residues are colored blue and red, and the helix and loop color are as per VMD default scheme. (B) Inter-residue wise contact maps for MSM states for glycosylated system.

**Figure 5.**
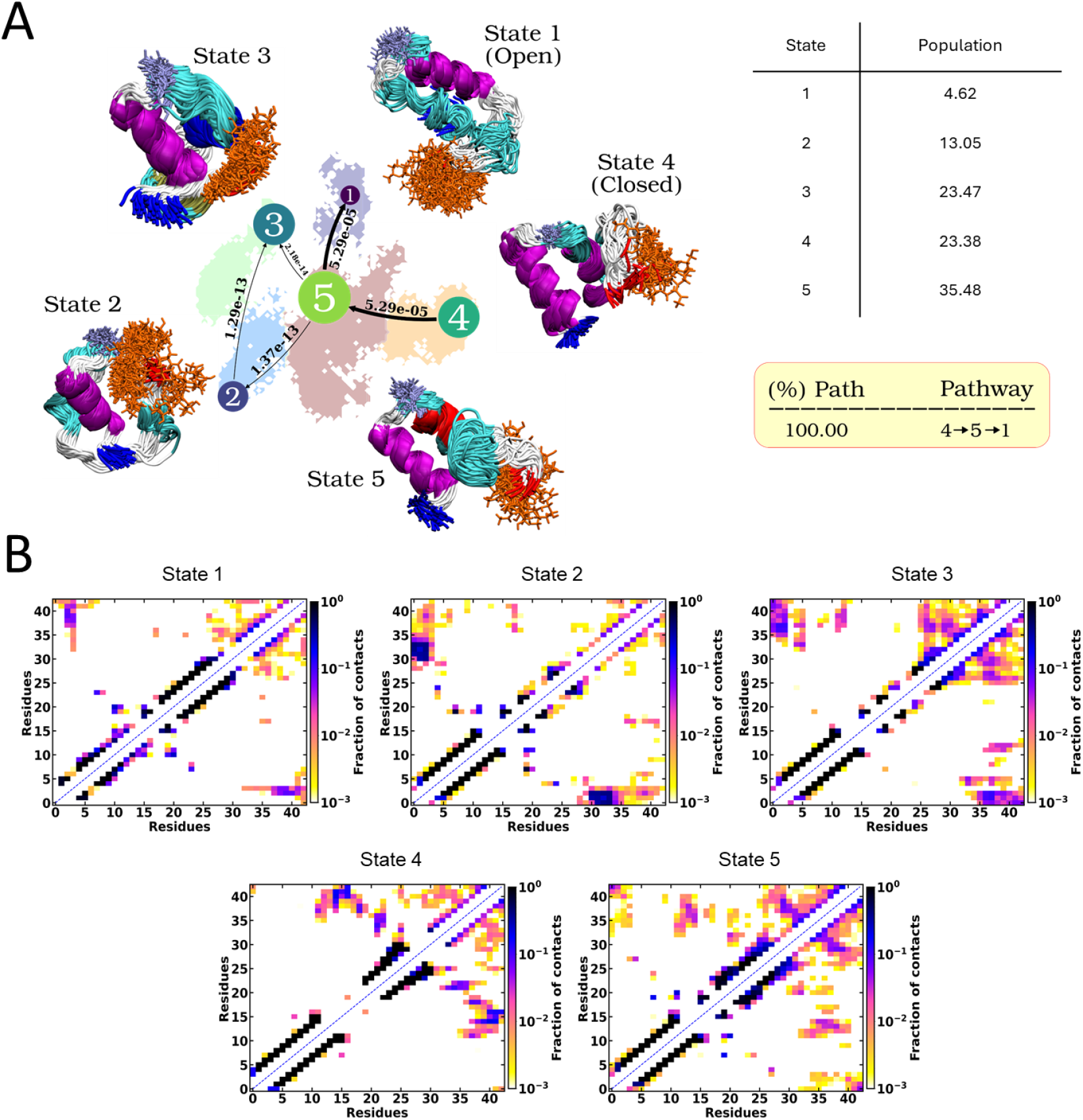
(A) Markov state models and the proportion of each state with unidirectional transition probability and pathways between states derived using latent space of Cosine Annealed VAE from methylated dataset. The N-terminal and C-terminal residues are colored blue and red, and the helix and loop color are as per VMD default scheme. (B) Inter-residue wise contact maps for MSM states for methylated system.

The α-methylated system’s MSM (Figure 5A) partitions the landscape into fewer states whose connectivity reflects preferred routes between loop-rigidified conformers, in agreement with residue-wise contact analysis (Figure 5B) and suppressed loop fluctuations (Figure 2B-C). Whereas, the metastates remain more interconnected with faster transition time in the *glycosylated* model, consistent with a stabilized yet somewhat more permissive landscape. The methylation metastable states revealed destabilization in one of the alpha-helix facilitated by interactions of both the two S/O-linked GlcNAc (Figure 4A & 5A).

In contrast, the *control* exhibits a more ramified kinetic graph (SI Figure S13) and state-resolved contact maps with fewer persistent contacts around the Ser18 loop (SI Figure S14), matching its broader FES and higher RMSF. Thus, establishing a progressive tightening of conformational space from the flexible *control* to the *glycosylated* and ultimately to the *methylated* form. (SI Table S1 & Figure 3).

Taken together, the stability profiles (Figure 2), FES topology (Figure 3; SI Figure S12), and MSMs (Figure 4A,5A & SI Figure S13) converge on a consistent mechanism: post-translational modifications that reinforce local packing around Ser18, either through glycan-mediated interactions or α-methylation-induced backbone restrictions that narrow the conformational and kinetic ensembles by stabilizing loop-packed conformers and elevating barriers to excursions into flexible states. This modification-dependent reshaping of the energy landscape provides a structural-kinetic basis for the reduced conformational plasticity observed in the *methylated* model relative to the *control* and *glycosylated* models (Figure 4A, 5A, SI Table S1). While glycosylation stabilizes the functional loop-helix coupling through organized yet interconvertible basins, α-methylation locks the backbone into sterically constrained geometries, distorting one α-helix as can be observed from the metastates 1-3 (Figure 5A) and kinetically trapping the peptide in non-functional conformations.

## 4 Discussion

Across three GccF variants, control (non-glycosylated), native glycosylated, and α-methyl-Ser18, we find that post-translational context around Ser18 is the dominant lever shaping the peptide’s conformational and kinetic organization. Glycosylation maintains the conformational plasticity by stabilizing the helical and inter-helical loop (residues ∼10–20), whereas α-methylation perturbs the native hydrogen-bonding network and restricts the conformational heterogeneity. These trends are concordant across stability metrics (RMSD/RMSF), latent-space free-energy landscapes, and MSM metastates and transition rates.

Latent-space FES from the cosine-annealed VAE consolidate this picture: native glycosylation expands the conformational ensemble into multiple well-defined basins that are rapidly interconnected, while α-methylation compresses the landscape into fewer, deeper minima, indicative of restricted flexibility and kinetic trapping within rigidified conformers. Together, these results indicate that α-methylation facilitated glycan-mediated packing around Ser18 kinetically “locks in” conformations and loss of conformational adaptability, while native glycosylation does not apply such steric constraints and maintains kinetic connectivity among functional conformers. Methodologically, the cosine-annealed VAE provided the most faithful dynamical embedding (highest cross-validated VAMP2 scores), allowing accurate reconstruction of state populations and transition pathways. The FES maps (SI Figure S12) and state-resolved contact networks (SI Figure S14; Figure 4B and 5B) collectively validate these mechanistic distinctions.

Overall, our MD-VAE-MSM framework reveals a modification-dependent reshaping of GccF’s conformational landscape: native glycosylation enforces structural order and kinetic persistence, whereas α-methylation at Ser18 disrupts native interactions and restricts conformational flux. These insights provide a structural-kinetic rationale for differences in functional behavior and suggest concrete design principles centered around Ser18-proximal packing that endeavor to preserve the native glycan-peptide contacts to tune activity in glycocin-like antimicrobial peptides.

## Authors contribution

*Conceptualization: A*.*S*.; *Data curation: D*.*K*.*S. and S*.*A*.; *Formal analysis: D*.*K*.*S*., *S*.*A. Investigation: D*.*K*.*S. (Simulations and Structural Analysis), S*.*A. (Simulations, VAE and MSMs); Methodology: D*.*K*.*S. (Simulations and Parametrization), Validation: A*.*S*.; *Writing-original draft: D*.*K*.*S*.; *Writing-review and editing: A*.*S*.

## Acknowledgement

*This work was supported by the Department of Atomic Energy, Government of India, under Project Identification No. RTI4007. D*.*K*.*S acknowledges Dr. Jagannath Mondal, Associate Professor, Tata Institute of Fundamental Research, Hyderabad, Telangana, India-500046 for providing the computing facilities*.

