## Supplementary material for "Leveraging Deep Learning and MD Simulations to Decipher the Molecular Basis of Attenuated Activity in Glycocin F": SI_GccF_ds_v5.pdf

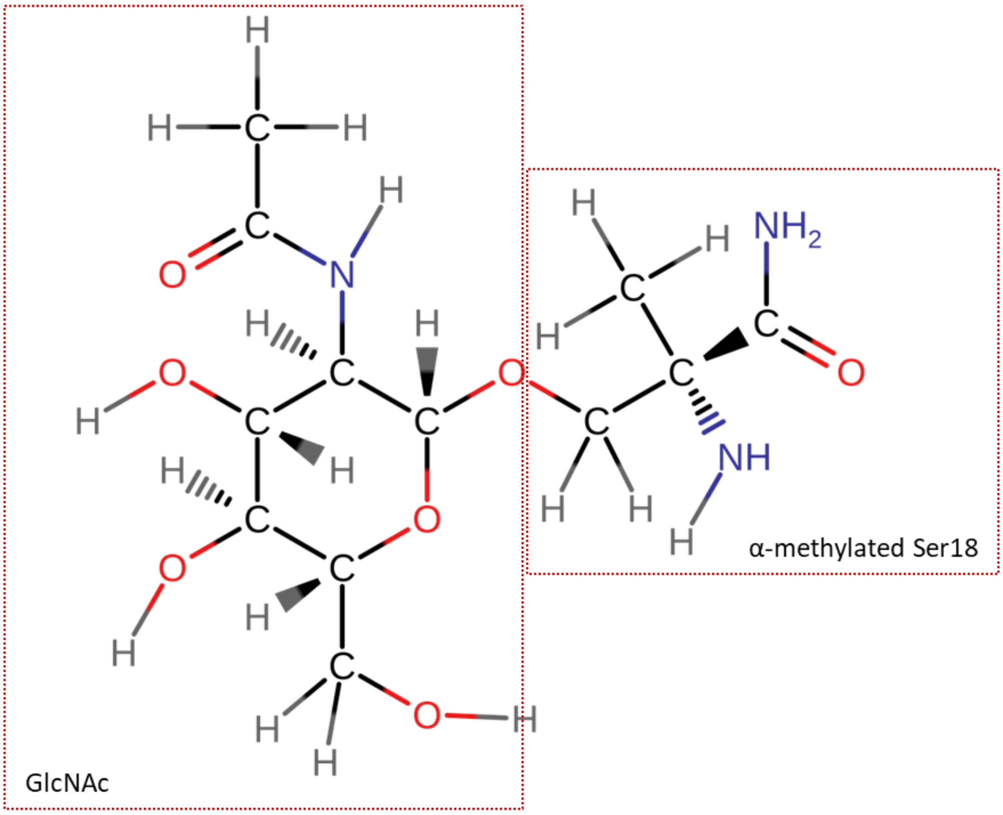


Figure S1. Schematic of N-Acetylglucosamine (GlcNAc) and α-methylated Ser18 used to generate force constants by the DFT Seminario method.

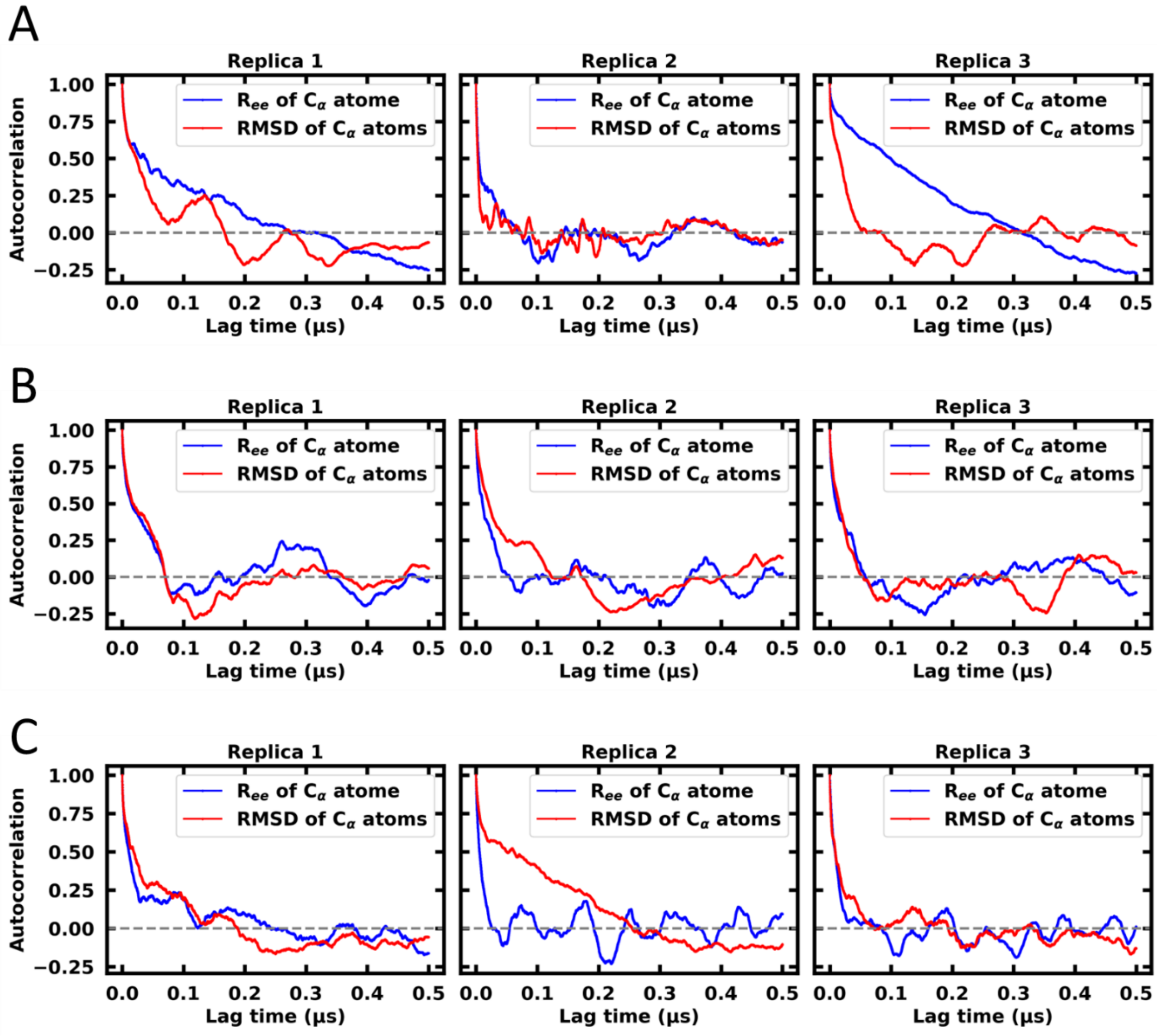


Figure S2. Convergence analysis with autocorrelation function for A. *control*, B. *glycosylated* and C. *methylated* systems. Time series of Cα end-to-end distance and Cα RMSD are depicted.

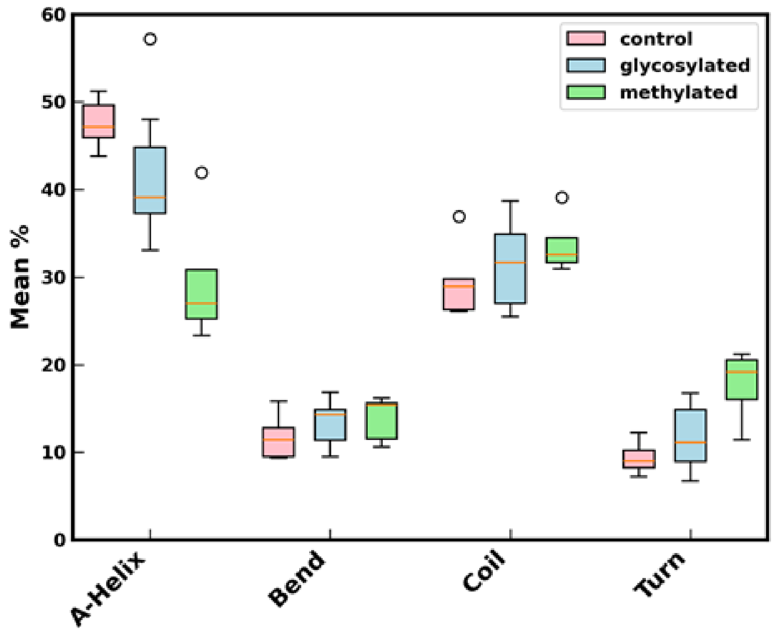


Figure S3. Percentage of secondary structure for *control*, *glycosylated* and *methylated* system.

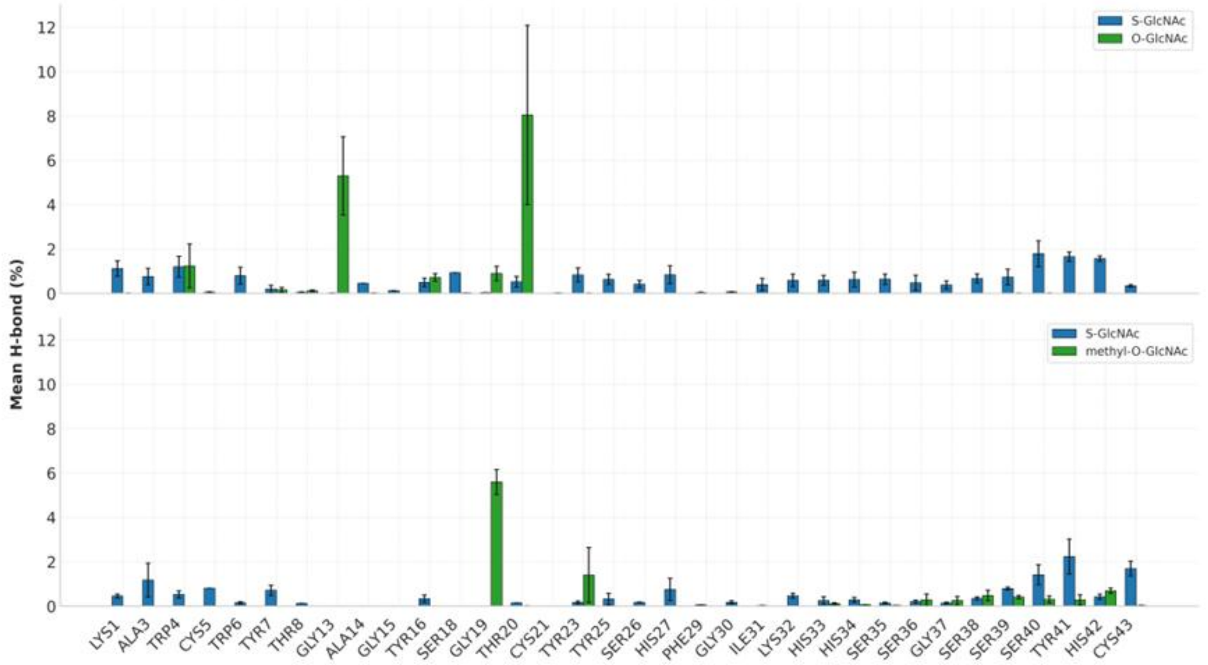


Figure S4. Average of hydrogen bond percentages calculated from three replicates simulations of *glycosylated* (top) and *methylated* (below) system. The S-linked and O-linked GlcNAc are shown in green and blue respectively.

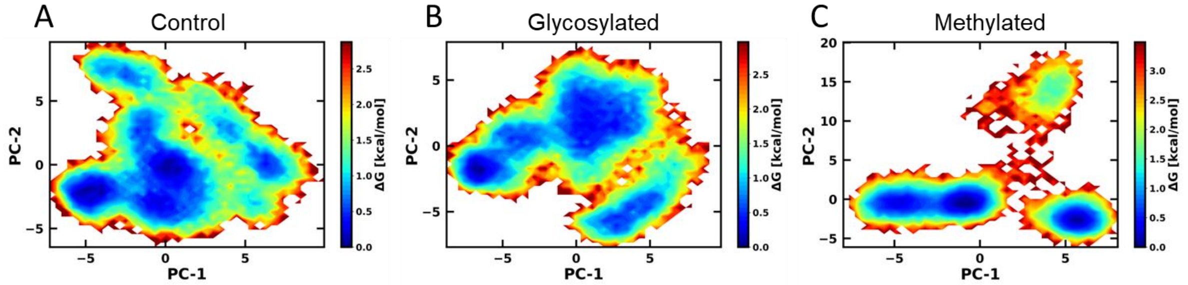


Figure S5. Free energy surface plotted of first two principal component from PCA of the dataset for A. *control*, B. *glycosylated* and C. *methylated* models.

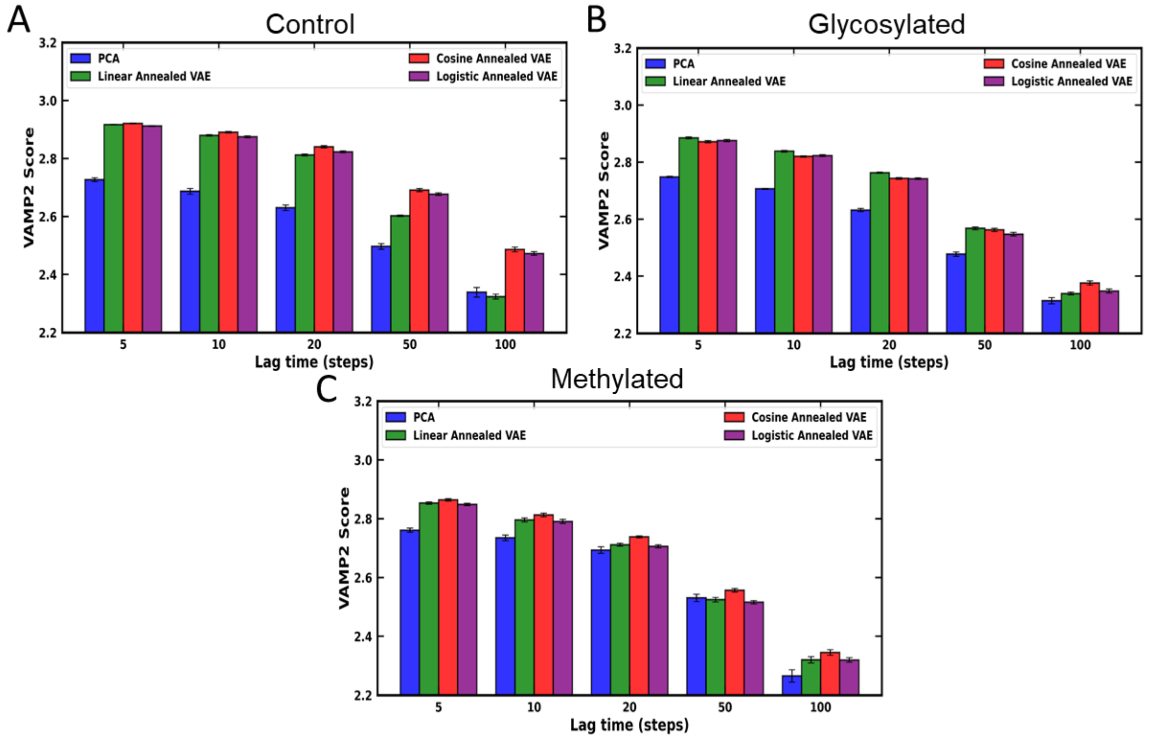


Figure S6. VAMP2 scores derived using PCA, Linear Annealed VAE, Cosine Annealed VAE and Logistic Annealed VAE method from the latent space projected from the simulation trajectories of A. *control*, B. *glycosylated* and C. *methylated* models.

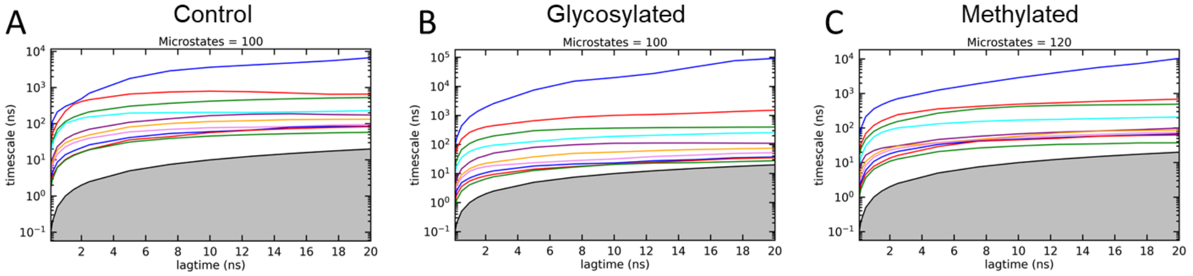

Figure S7. Implied time scale (ITS) depicting the slow evolving processes calculated considering ~100 microstates for A. *control*, B. *glycosylated* and C. *methylated* simulation data.

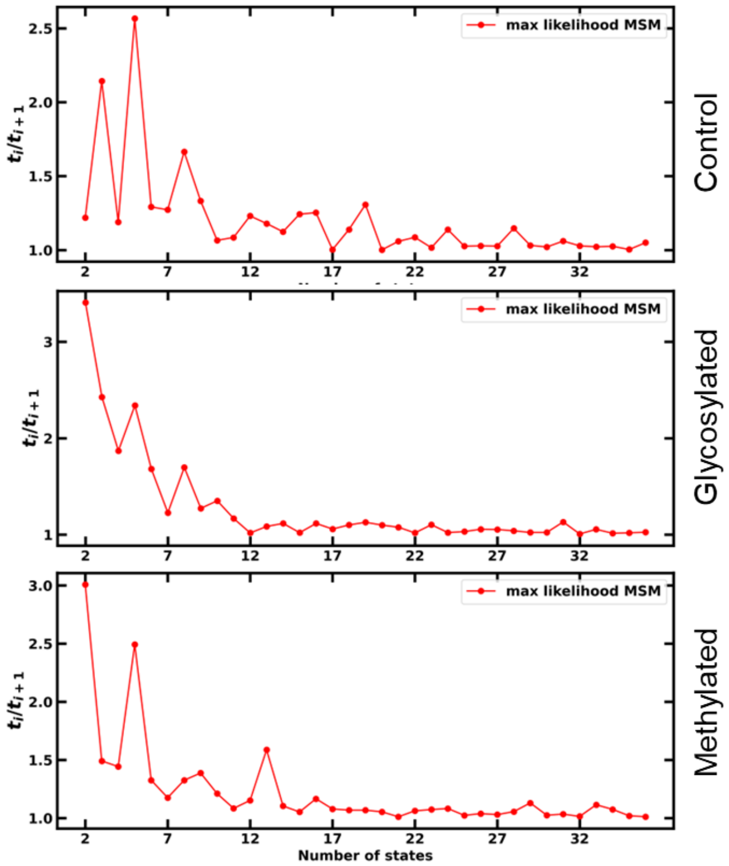


Figure S8. Spectral analysis depicting the separation of spectral states for *control*, *glycosylated* and *methylated* projected data.

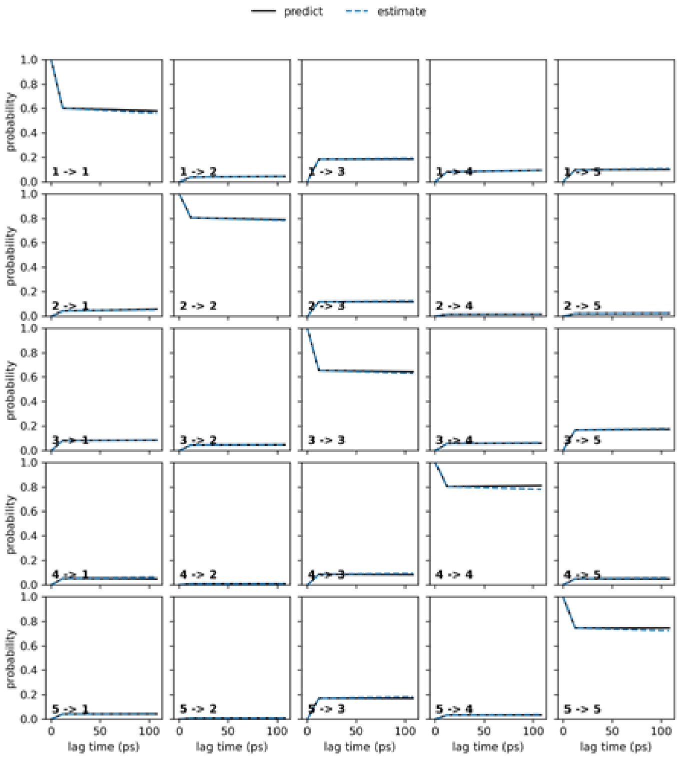


Figure S9. Chapman-Kolmogorov test predicted and estimated value of resolved states of *control* simulations shown for lag time of 100 ps.

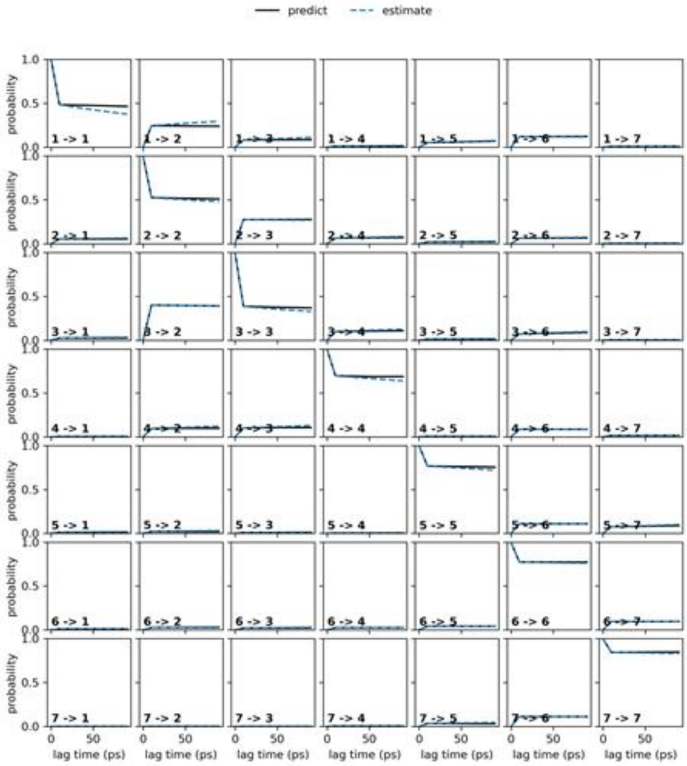


Figure S10. Chapman-Kolmogorov test predicted and estimated value of resolved states of *glycosylated* simulations shown for lag time of 100 ps.

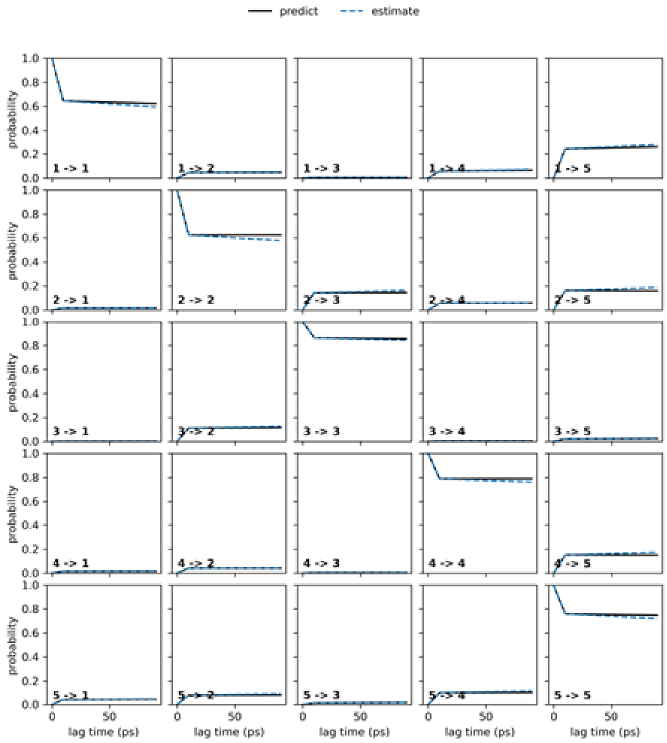


Figure S11. Chapman-Kolmogorov test predicted and estimated value of resolved states of *methylated* simulations shown for lag time of 100 ps.

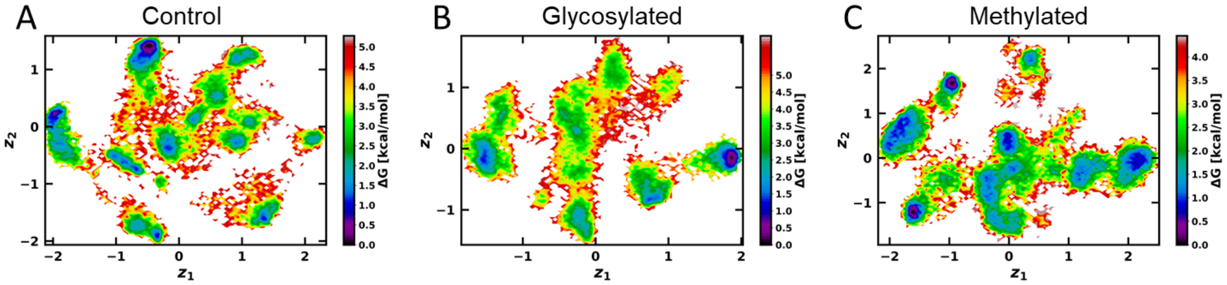


Figure S12. Free energy surface plotted on the latent space obtained by β-annealed VAE method for A. *control*, B. *glycosylated* and C. *methylated* models.

Table S1. Transition time (μs) of metastable states from "closed" to "open" obtained for studied models.

| *Control* | |
|---|---|
| **States** | **closed** to **open (μs)** |
| 1* | 0.82 |
| 2 (closed) | |
| 3* | 0.33 |
| 4* | 1.21 |
| 5 (open) | 0.51 |
| *Glycosylated* | |
| 1* | 0.46 |
| 2* | 0.02 |
| 3* | 0.06 |
| 4 (closed) | |
| 5* | 0.78 |
| 6* | 0.90 |
| 7 (open) | 1.63 |
| *Methylated* | |
| 1 (open) | 1.52 |
| 2* | 0.52 |
| 3* | 1.10 |
| 4 (closed) | |
| 5* | 0.09 |
| *Source = open; Sink = closed; *Treated as Source* | |

**Control**

| State | Population |
|---|---|
| 1 | 8.35 |
| 2 | 10.84 |
| 3 | 39.15 |
| 4 | 17.36 |
| 5 | 24.30 |

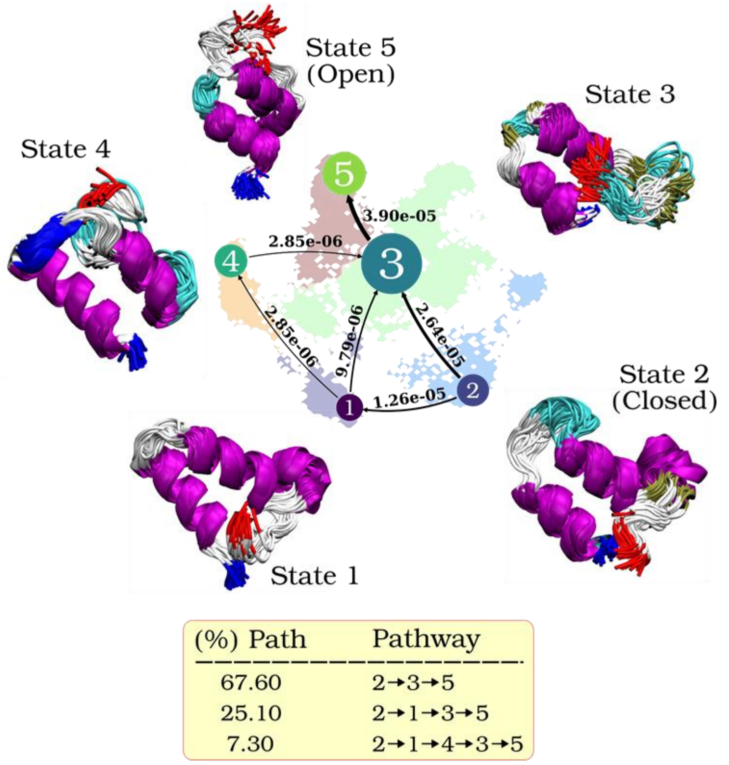


Figure S13. Markov state models and the proportion of each state with unidirectional transition probability and pathways between states derived using latent space of Cosine Annealed VAE from *control* dataset. The N-terminal and C-terminal residues are colored blue and red, and the helix and loop color are as per VMD default scheme.

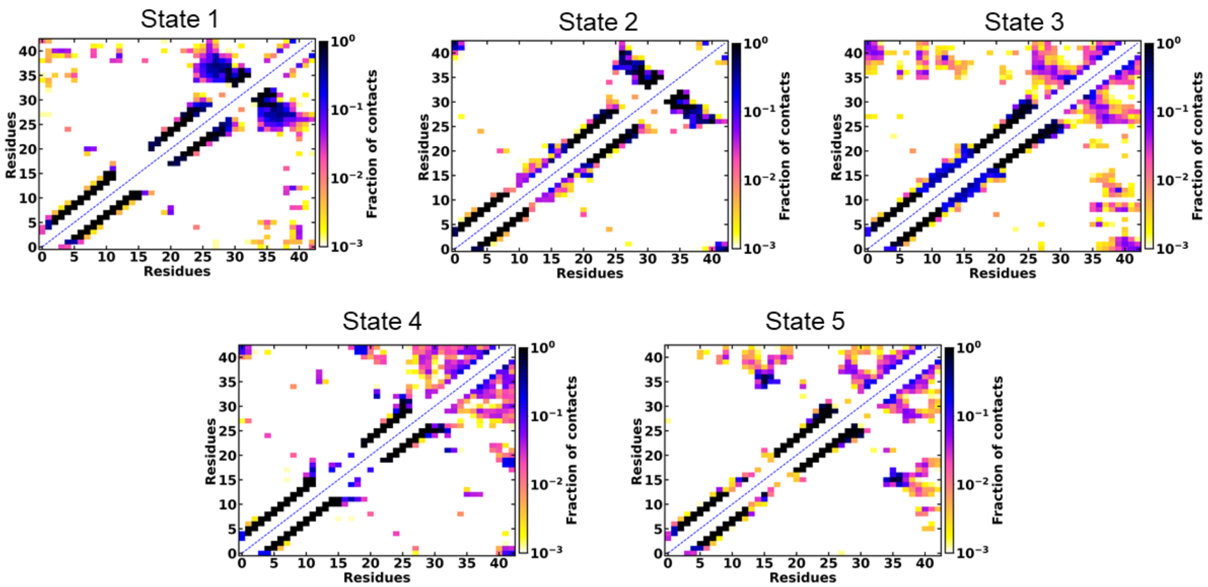


Figure S14. Intra-residue wise contact map and fraction of contacts for resolved MSMs of *control* dataset.
